# Discovering a new biocatalytic potential of milk fat globules: *in situ* conversion of exogenous docosahexaenoic acid to its oxylipins

**DOI:** 10.64898/2026.05.26.727978

**Authors:** Tana Hernandez-Barrueta, Nitin Nitin, Ameer Y. Taha

## Abstract

Milk fat globules (MFG) are complex structures that emulsify the fat in milk and exhibit various biological activities. Using docosahexaenoic acid (DHA) as a model polyunsaturated fatty acid, we demonstrate for the first time that a MFG-enriched fraction (isolated from commercial raw and unhomogenized bovine milk) catalyzes the oxidation of exogenous polyunsaturated fatty acid to its oxylipins. Static incubation of the MFG-enriched fraction (at 20% w/v in phosphate buffer and 5% v/v ethanol) with 150 µM of DHA for 1 h resulted in the production of ∼13 pmol/mg cream of DHA-derived oxylipins in free and esterified (i.e., bound) forms. High enrichment in free DHA-oxylipins was observed, wherein incubation with DHA resulted in >80% of all free oxylipins being derived from DHA, in contrast to control samples in which only <8% of all free oxylipins were DHA derivatives. The most abundant products generated were 17-hydroxydocosahexaenoic acid and 19(20)-epoxydocosapentaenoic acid. Oxylipin generation was dependent on the structural integrity of MFG, with mechanical disruption (vortexing) impairing oxylipin synthesis more severely than thermal treatment. These findings suggest that MFGs harbor multiple metabolically active enzymes, including cytochrome P450, lipoxygenases, acyl-CoA synthetases, and acyltransferases, that act cooperatively to synthesize and esterify DHA-derived oxylipins. In summary, this study highlights the dual potential of MFGs to serve both as a food-grade biocatalyst for the synthesis of DHA-derived oxylipins and as carriers for these bioactive lipids. Future studies should evaluate the bioavailability and physiological relevance of these metabolites when consumed in the diet.

## 1. Introduction

The anti-inflammatory properties associated with consumption of polyunsaturated fatty acids (PUFAs) are in part mediated by their oxidized metabolites, known as oxylipins [reviewed in ^1^]. Oxylipins are produced by cyclooxygenase, lipoxygenase (LOX), or cytochrome P450 (CPY450) enzymes with ω-hydroxylase or epoxygenase activity, or through nonenzymatic auto-oxidation [reviewed in ^1^]. In particular, oxylipins derived from the omega-3 fatty acids docosahexaenoic acid (DHA) and eicosapentaenoic acid (EPA) are recognized for their role in regulating the resolution of inflammation in different tissues ^2–4^. Circulating and tissue concentrations of these omega-3-derived oxylipins are often dysregulated in chronic inflammatory conditions such as diabetes, ^5^ obesity, ^6^ and cardiovascular disease.^7^ As a result, supplementation with omega-3 PUFAs has been proposed to increase endogenous oxylipin levels. However, DHA turnover is relatively slow. For instance, a study in healthy adults showed that 3 months of supplementation with DHA was required to double circulating levels of DHA-derived epoxides relative to baseline, and extending supplementation for up to 12 months (even at higher DHA doses) did not further increase the levels of these oxylipins.^8^ Moreover, the efficiency of DHA and EPA conversion to oxylipins is influenced by gender ^9^ and genetic factors ^10,11^. Therefore, rather than relying solely on dietary DHA supplementation, an alternative approach might be to generate and deliver the omega-3 oxylipins themselves.

Oxylipins can be chemically synthesized, as shown for epoxide and hydroxy derivatives of DHA and EPA ^12–14^, enabling the study of their biological effects in animal models. However, this approach is costly and unsustainable for large-scale use due to the generation of chemical waste and extensive purification steps. More sustainable alternatives include fermentation, such as Rorrer et al.’s use of seaweed cell cultures to produce hydroxylated PUFAs via native LOX pathways ^15^. Microbial enzymes are another option: CYP450 from *Bacillus megaterium* (expressed in *E. coli*) and fungal peroxygenases have been used to convert DHA to its epoxides ^16,17^. However, because pure enzymes may be commercially limited or prohibitively expensive, a promising alternative is to use food ingredients that may have the enzymatic capacity to convert DHA into oxylipins. Still, both these chemical and enzymatic approaches may face challenges with the stability of the generated oxylipins during storage, processing (e.g., cooking), and oral delivery. For instance, the bioavailability of a mono-hydroxy derivative of linoleic acid has been demonstrated in rats ^18^, but this cannot be assumed for structurally different oxylipins like epoxides, which can undergo ring-opening in acidic environments ^19^. Altogether, this highlights the need for identifying food-based systems capable of simultaneously generating and stabilizing oxylipins.

Milk fat globules (MFGs) are micrometer-size structures that emulsify the fat in milk. They consist of a core of neutral lipids enclosed by a tri-layer phospholipid membrane known as the milk fat globule membrane (MFGM) ^20^. The MFGM features various enzymes, including some of which retain their activity in milk, the most well-documented being xanthine oxidase ^21,22^. These enzymes are incorporated into the MFGM during MFG synthesis in mammary epithelial cells, as the MFGM originates from the cytoplasmic and rough endoplasmic reticulum membranes of these cells (reviewed by ^20,23,24^). Given that mammary epithelial cells express enzymes such as CYP450 ^25,26^, which are known to convert PUFAS to their oxylipins, it is reasonable to hypothesize that MFGs may inherit this biocatalytic potential. Additionally, the endogenous presence of oxylipins in MFGs (albeit those derived from DHA and EPA constitute <5%) ^27,28^ suggests an inherent potential to serve as carriers for these compounds. In this context, we hypothesized that MFGs, due to their enzymatic potential and structural complexity could serve as both biocatalysts and encapsulation matrices for PUFA-derived oxylipins. We therefore investigated an approach in which precursor PUFAs are loaded into MFGs to enable their putative enzymatic transformation, using our previously published method to encapsulate hydrophobic compounds into intact MFGs via passive infusion ^29^.

Thus, this study aimed to evaluate the potential of MFGs to catalyze the conversion of PUFAs to their oxylipins, using DHA as a model PUFA due to the clinical relevance of its oxylipins. A MFG-enriched fraction was obtained from raw, unhomogenized commercial bovine milk to ensure the inclusion of intact globules representative of multiple animals and to preserve potential enzymatic activity. This fraction was incubated with or without exogenous DHA, and the resulting oxylipins were quantified. Given that milk contains oxylipins in both free (i.e., unbound) and esterified (i.e., bound) forms ^27^, we employed our previously validated protocols to extract and analyze free and total oxylipins (i.e., free + esterified) using liquid chromatography coupled to tandem mass spectroscopy (LC-MS/MS) ^27,28^. Finally, we evaluated the role of MFG integrity in oxylipin generation by subjecting the globules to mechanical and thermal treatments.

## 2. Materials and methods

### 2.1. Materials

A commercial brand of raw, unhomogenized, grade A milk obtained from Jersey cows (Claravale Farm, CA) was employed in this study. For each experiment, milk was acquired from a local store (Davis Food Co-op, Davis California) and kept at 4 °C until use, no later than 2 days after its purchase.

Phosphate buffer solution 10X (PBS, Cat # BP3994), methanol (Cat #A454-4), LC/MS grade methanol (CAS #67-56-1), ethyl acetate (Cat #E196-4), chloroform HPLC grade (Cat #C607-4) and Oregon green™ 488 DHPE (Cat #O12650) were purchased from Thermo Fisher Scientific. Triphenylphosphine (TPP; Cat #T84409-100G), butylated hydroxytoluene (BHT; Cat #W218405-SMPLE-K), ethylenediaminetetraacetic acid (EDTA, Cat #EDS-100G), acetic acid (Cat # 695092-100ML), sodium chloride (NaCl, Cat # S7653-250G), EDTA-2Na (Cat #E5134-50g) and sodium hydroxide (NaOH; Cat # S5881-500G) were acquired from Sigma-Aldrich. Ethanol (Cat #UN1170) was from Koptec.

Docosahexaenoic acid (DHA, Cat # 90310), unlabeled oxylipin standards used to generate the calibration curve and deuterated oxylipin standards used as surrogates during oxylipin extraction were from Cayman Chemical (Ann Arbor, MI, USA). The surrogate standards were d11-11,12-epoxyeicosatrienoic acid (EpETrE, Cat # 10006413), d11-14,15-dihydroxyeicosatrienoic acid (DiHETrE, Cat #1008040), d4-6-keto-prostaglandin (PGF1α, Cat #315210), d4-9-hydroxyoctadecadienoic acid (HODE, Cat # 338410), d4-leukotriene B4 (LTB4, Cat #320110), d4-prostaglandin E2 (PGE2, Cat #314010), d4-thromboxane B2 (TXB2, Cat #319030), d6-20-hydroxyeicosatetraenoic acid (HETE, Cat #390030) and d8-5-hydroxyeicosatetraenoicacid (HETE, Cat # 334230).

### 2.2. Methods

#### 2.2.1. Isolation of milk fat globules

Milk fat globules were isolated from milk as described by Alshehab et al.^30^, with slight modifications. Briefly, milk was diluted in PBS 1X (prepared with MilliQ water) at a 1:9 ratio, and centrifuged at 3,000 × g for 5 minutes at 4°C. The cream layer was collected, resuspended at 4% (w/v) in PBS 1X, and centrifuged under the same conditions. The cream recovered from this step was used for the following experiments and will be referred to as ‘MFG-enriched fraction’.

#### 2.2.2. Incubation of MFG-enriched fraction with DHA

One gram of MFG-enriched fraction was suspended in a mixture of 4.75 mL of PBS 1X and 100 µL of ethanol, to which 150 µL of an ethanolic solution of 5 mM DHA was spiked (final concentrations: MFG-enriched fraction 20% w/v, ethanol 5% v/v, and DHA 150 µM). To account for oxylipins naturally found in the milk fraction, negative controls without exogenous DHA were considered, and to account for oxylipins possibly found in the DHA stock, a control DHA stock was prepared at the same final concentration in PBS and ethanol as the samples. Experiments were performed in quadruplicates per group (unless otherwise specified).

Tubes were gently mixed by inversion and allowed to incubate statically for 60 minutes at room temperature in the dark. Aliquots of 600 µL were taken out from the same tube (i.e., repeated measurement) 0 and 60 minutes after addition of the DHA ethanolic solution (or equivalent volume of ethanol for the controls). The aliquots were immediately centrifuged at 3,000 × g for 5 min at 4 °C and the milk cream layer was weighted, collected and stored at -80 °C for further analysis. Note that because of this, there was a ∼10 minutes delay between sampling and freezing due to the centrifugation and weighting of the sample prior to placing them in the freezer.

To evaluate the role of globule integrity on the generation of DHA-derived oxylipins, two approaches were used: mechanical and thermal treatment. For the mechanical treatment, the MFG-enriched fraction was diluted in PBS and ethanol, and the DHA solution was added (n= 3 per treatment). All final concentrations and incubation conditions were as described before, except that in these samples, tubes were vortexed after the addition of DHA (or ethanol for controls) and immediately before taking an aliquot at 0, 15, 30, 45 and 60 minutes. For the thermal treatment, the MFG-enriched fraction was first diluted at 20 % w/v, heated at 90 °C for 10 minutes, and then cooled down to room temperature before adding the ethanol and DHA solution (or ethanol for control samples). All final concentrations, incubation conditions and sampling procedures were as described before.

#### 2.2.3. Imaging of the MFGs and particle size distribution analysis

To assess the structural integrity of the MFGs after centrifugal separation, a triplicate of MFG-enriched samples was independently prepared (as described in section 2.2.1.) and diluted to 20% w/v in PBS 1X before analysis. Two aliquots of 40 µL of the MFG-enriched fraction were suspended in 160 µL of PBS 1X. One aliquot was stained with 1.5 µL of Oregon Green 488 DHPE (1 mg in 100 µL of chloroform) to probe the MFG membrane ^30^. The other was stained with 1 µL of Nile Red (1 mg/mL in acetone) to probe the core of the MFG, as this dye stains neutral lipids ^31,32^. Samples were then left to incubate in the dark for 10 minutes. Fluorescence images were collected using an inverted optical microscope (Olympus IX-7, Olympus Inc., Center Valley, PA, USA) with a 40X oil immersion objective (Olympus UPlanFL, 40×/1.30 oil, Olympus Inc., Center Valley, PA, USA). An excitation filter 480/30 nm and an emission filter 570/60 nm were used. The remaining samples (after removing the aliquots for imaging) were diluted 1:1 volume ratio in 35 mM of EDTA at pH 7. The particle size distribution was quantified using a Microtrac S3500 (Microtrac instrument, US) with the following parameters: 1.46 and 1.33 refractive index of milk and water, respectively; absorbance 0.001. Raw milk (i.e., before centrifugal separation) was also analyzed as a control.

To evaluate the role of globule integrity on the generation of oxylipins, at the end of the 60 minutes incubation of the vortexed and preheated MFG-enriched fraction with DHA, extra 40 µL aliquots were taken and stained with Nile Red as described above.

#### 2.2.4. Extraction of oxylipins

For the extraction of free and total (i.e. free + esterified) oxylipins, a mixture of surrogate standards solution was prepared such that each of the following compounds was at a final concentration of 2 μM in methanol (LC/MS grade): d11-11(12)-EpETrE, d11-14,15-DiHETrE, d4-6-keto-PGF1a, d4-9-HODE, d4-LTB4, d4-PGE2, d4-TXB2, d6-20-HETE, and d8-5-HETE. Other solutions used were 1) antioxidant solution containing TPP, BHT, and EDTA each at 0.2 mg/mL in MilliQ/methanol 1:1 v/v, filtered through a 0.45 μm Millipore filter, 2) extraction solvent encompassing methanol with 0.1% acetic acid and 0.1% BHT and 3) solid phase extraction buffer made of 0.1% acetic acid and 5% methanol in MilliQ water. The extraction of free and total oxylipins was performed according to Teixeira et al. ^28^, with minor modifications as described below.

##### 2.2.4.1. Extraction of free oxylipins

Approximately 10 mg of milk cream was resuspended in 190 μL of 1X PBS, to which 600 μL of precooled methanol, 200 μL of extraction solvent, 10 μL of antioxidant solution, and 10 μL of surrogate standard solution were added. Samples were mixed by vortexing and centrifuged at 15,000 × g for 10 min at 0 °C. The supernatant was collected (∼0.85 mL) and MilliQ water was added to adjust the methanol concentration to ∼15% v/v (∼4.5 mL). Samples then underwent solid phase extraction.

##### 2.2.4.2. Extraction of total oxylipins

Approximately 10 mg of milk cream were subjected to Folch extraction with 3 mL of chloroform/methanol (2:1 v/v) containing 0.002% BHT, and 0.75 mL of 0.9% NaCl with 1 mM EDTA-2Na. Samples were vortexed and centrifuged at 598 × g for 10 min at 0 °C. The organic phase was collected. To the aqueous phase, 2 mL of chloroform was added, followed by vortexing and centrifugation at 598 × g for 10 min at 0 °C. The organic phases (containing the total lipid extract) were combined and dried under N_2_, reconstituted in chloroform/isopropyl alcohol (2:1 v/v), and 1 mL was dried again under N_2_. To this, 200 μL of extraction buffer, 10 μL of antioxidant solution, and 10 μL of surrogate standard solution were added. Samples were hydrolyzed at 60 °C for 30 min with 200 μL of NaOH 0.4 M in MilliQ water:methanol (1:1 v/v) and allowed to cool down for 5 minutes. Twenty-five μL of acetic acid were then added to acidify samples to pH 4-6 (confirmed in a few representative samples using litmus paper), followed by the addition of 1575 μL of MilliQ. Samples then underwent solid phase extraction.

##### 2.2.4.3. Solid phase extraction

Total and free oxylipins were purified using 60 mg Oasis HLB columns (3 cc, Waters Corporation, CA, USA, Cat# WAT094226) prewashed with one column volume of ethyl acetate followed by two of methanol and one of solid phase extraction buffer. Samples were poured onto the column, rinsed with two volumes of solid extraction buffer, and dried under vacuum (∼15 psi, 20 minutes). Oxylipins were eluted with 0.5 mL of methanol and 1.5 mL of ethyl acetate, dried under N_2_, reconstituted on 100 μL of methanol (LC/MS grade), and filtered in a Ultrafree-MC centrifugal filters (0.1μm; Millipore Merck, Burling-ton, MA, USA, #UFC30VV00) centrifuged at 15000 × g for 2 min at °C. Samples were stored at -80 °C until analyzed by ultra-high performance liquid chromatography coupled to tandem mass spectrometry (UPLC-MC/MS).

#### 2.2.5. UPLC-MS/MS analysis

A total of 73 oxylipins were quantified using a 1290 Infinity ultra-high-pressure-liquid chromatography (UHPLC) system coupled to a 6460 triple-quadrupole tandem mass spectrometer (MS/MS) with electrospray ionization (Agilent Technologies, Santa Clara, CA, USA). Oxylipins were separated using a ZORBAX Eclipse Plus C18 column (2.1 mm by 150 mm; particle size 1.8μm; Agilent Technologies, Santa Clara, CA, USA). Mobile phase A encompassed 0.1% acetic acid in MilliQ water; mobile phase B was 0.1% of acetic acid in acetonitrile:methanol (85:15 v/v). The total run time was 20 min as follows. The flow gradient started with mobile phase A at 65%, and subsequently changed to 15, 0, and 65% at minutes 12, 15, and 17 respectively. During this time, the flow rate was 0.25 mL/min from 0 to 15 min, 0.4 mL/min from 15 to 19 min, and 0.30 mL/min from 17 to 20 min. Auto-sampler and column were maintained at 4 and 45 °C, respectively. The drying gas temperature was 300 °C with a flow of 10 L/min, sheath gas temperature was 350 °C at a flow of 11 L/min, and the nebulizer pressure was 35 psi.

The samples were injected at a volume of 10 μl, analyzed in negative electrospray ionization mode, and captured using optimized dynamic multiple reaction monitoring. Details on the optimized method (precursor and product ions, fragmentor, multiple reaction monitoring, collision energies and retention times) for each analyte have been reported before ^28^ and are summarized in Supplementary Table S1 next to the limits of detection.

### 2.3. Statistical analysis

The particle size distribution of MFGs from raw milk and the MFG-enriched fraction was compared via unpaired t-test. Two-way repeated measures analysis of variance (ANOVA) was conducted to determine the effect of addition of exogenous DHA and incubation time on the concentration of total DHA-oxylipins in the MFG-enriched fraction (separately performed for the MFG-enriched fraction, the preheated MFG-enriched fraction and the vortexed MFG-enriched fraction); the p-values from these analyses can be found in Supplementary Table S2. ANOVA was not performed for free DHA-oxylipins since these were mostly non detected in all replicates of treatments without addition of exogenous DHA. All statistical analyses were performed using R (version 4.2.1).

## 3. Results

### 3.1. Isolation and characterization of MFGs from raw unhomogenized milk

A MFG-enriched fraction was obtained from raw unhomogenized bovine milk by two steps of centrifugal separation. The structural integrity of the MFGs in the enriched fraction was evaluated by measuring the particle size distribution, and by imaging the neutral lipids in the MFG core using Nile Red, and the MFGM using the phospholipid-labeled fluorescent probe Oregon Green™ 488 DHPE ( Figure ***1***).

**Figure 1.**
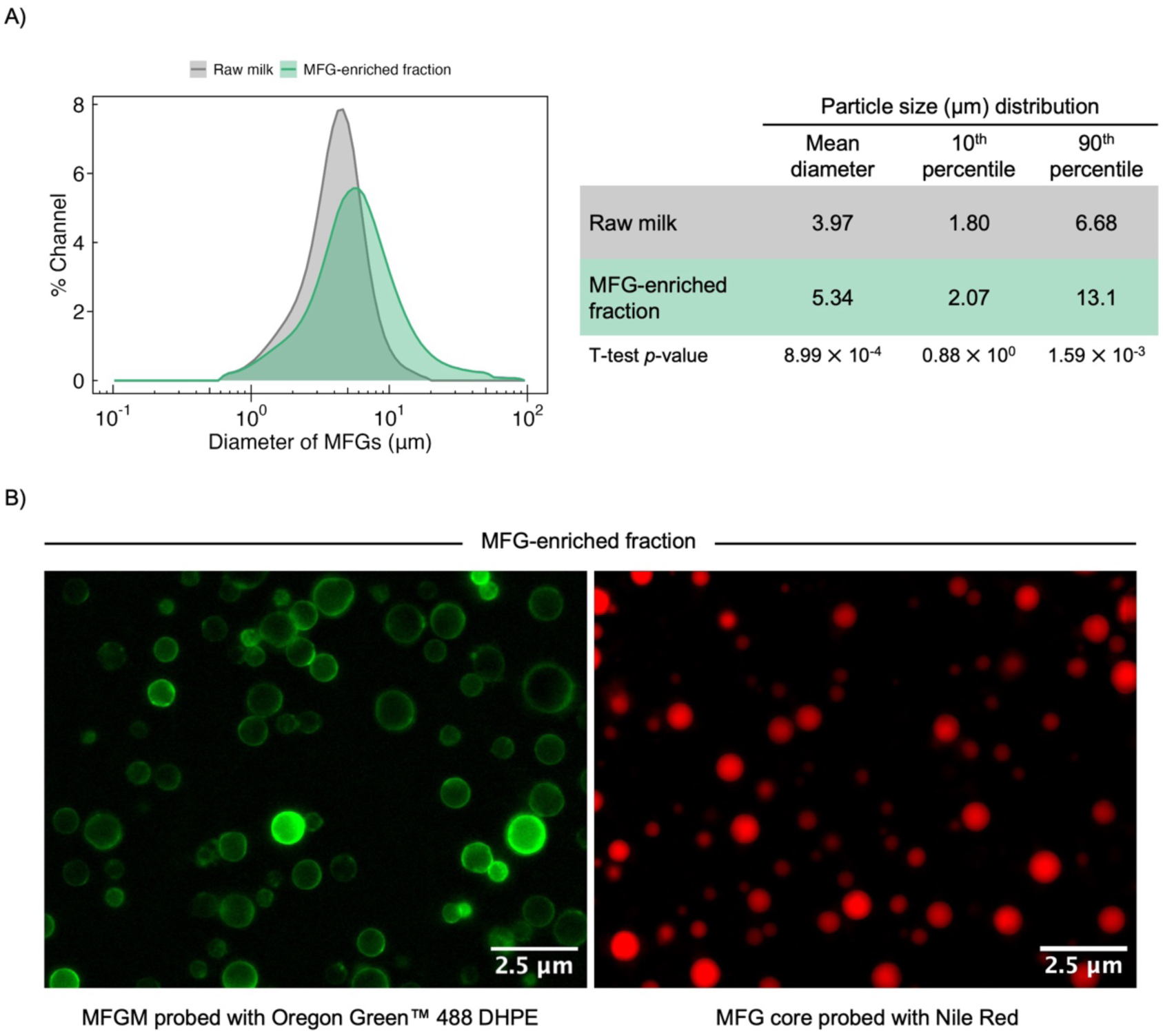
Morphological characterization of raw unhomogenized bovine milk and its milk fat globule (MFG) enriched fraction obtained by 2 cycles of centrifugal washing. A) Particle size distribution. Results in the table are presented as mean ± standard deviation of n=3. B) Fluorescence images of MFG-enriched fraction.

Compared to raw milk, MFGs in the enriched fraction exhibited significantly higher mean and 90th percentile diameters, approximately 1.3- and 2-fold greater, respectively (Figure 1A). This increase in particle size is attributed to MFG coagulation during enrichment, rather than disruption of the MFGs, as confirmed by the preserved membrane and core structure observed in the fluorescent micrographs (Figure 1B).

### 3.2. Oxylipins are generated in MFG-enriched fraction upon incubation with exogenous DHA

Having confirmed the structural integrity of the MFGs from the MFG-enriched fraction, the latter was incubated for 60 minutes at room temperature with or without the addition of exogenous DHA (150 µM). After incubation, the oxylipins were extracted and quantified using LC-MS/MS. **Table 1** shows the resulting concentration of DHA-derived oxylipins, while concentrations of other PUFA-derived oxylipins are presented in Supplementary Table S3.

**Table 1.**
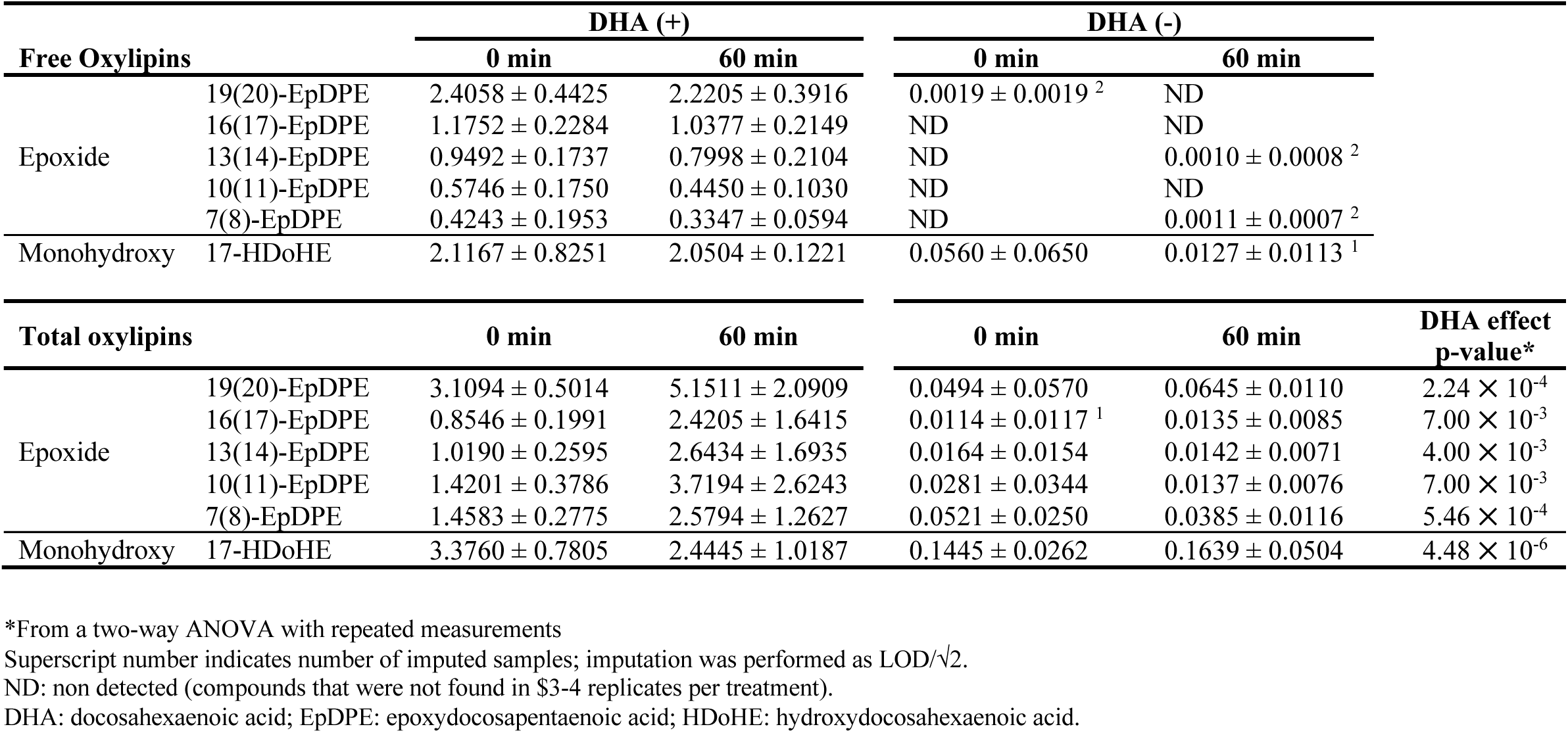
Concentration of DHA-derived oxylipins (pmol/mg of cream) in MFG-enriched fraction incubated at room temperature during 1 h in the presence (+) or absence (-) of exogenous DHA (150 µM). Results are reported as the average ± standard deviation (n=4).

In the absence of exogenous DHA, most free DHA-derived oxylipins were undetectable or present at trace levels (< 0.05 pmol/mg). In contrast, 60 minutes of incubation with exogenous DHA led to markedly higher concentrations: free DHA-epoxides (epoxydocosapentaenoic acid, EpDPE) ranged from 0.3 to 2.4 pmol/mg cream depending on the regioisomers, and free DHA-mono-hydroxy (hydroxydocosahexaenoic acid, HDoHE) concentrations were 2.0 to 3.5 pmol/mg cream. Two-way repeated measures ANOVA could not be performed for free DHA-oxylipins due to all the non-detected values in samples without addition of DHA.

In the total (i.e., free + esterified) oxylipin pool, two-way repeated measures ANOVA revealed that incubation with exogenous DHA—but not incubation time—significantly (*p* < 0.01) affected the concentration of EpDPEs and HDoHE. When no DHA was added, total DHA-oxylipins were detected below 0.15 pmol/mg of cream, while the addition of exogenous DHA resulted in concentrations of total EpDPEs and HDOHE of 0.8 to 5.1 and 2.4 to 3.7 pmol/mg of cream, respectively. Although aliquots were taken after 0 and 60 minutes of incubation, in each case, there was an extra ∼10 minutes of contact between DHA and the MFGs as the aliquots were centrifuged and weighted before being stored at -80 °C (as described in section 2.2.2). Therefore, the relatively high concentration of DHA-oxylipins in samples incubated with DHA for ‘0 minutes’ reflects this delay in freezing, rather than high baseline levels (for example, if oxylipins preexisting in the DHA stock had partitioned into the MFGs). This was confirmed as the DHA stock showed substantially lower concentrations of DHA-oxylipins <0.3 pmol/mg (Supplementary Table S4) compared to when incubated in the presence of MFG-enriched fraction.

Taken together, our results indicate that the addition of exogenous DHA to the MFG-enriched fraction led to the formation of DHA-derived EpDPEs and HDoHE, resulting in concentrations one to two orders of magnitude higher than those observed in control samples. Among the DHA-oxylipins detected, 19(20)-EpDPE and 17-HDoHE were the most abundant species. DHA diols (dihydroxypentaenoic acid) were non-detected or present at very low concentrations (< 0.01 pmol/mg cream) regardless of the addition of DHA to the MFG-enriched fraction, suggesting minimal generation during the 60-minute incubation period (see Supplementary Table S3).

In **Table 1**, a trend was observed wherein the concentration of total DHA-epoxides in samples spiked with DHA increased after 60 minutes of incubation, while the free epoxide pool remained relatively unchanged over time. To explore this further, we estimated the esterified pool by subtracting the concentration of free from the concentration of total oxylipins. As illustrated in **Figure 2**, the relative ratio of esterified to free EpDPEs increased over time (from 22% to 57% for 19(20)-EpDPE, 0% to 57% for 16(17)-EpDPE, 7% to 70% for 13(14)-EpDPE, 60% to 80% for 10(11)-EpDPE, and 70% to 87% for 7(8)- EpDPE). This suggests that a continuous esterification of DHA-derived epoxides occurred during the incubation of the MFG-enriched fraction with exogenous DHA.

**Figure 2.**
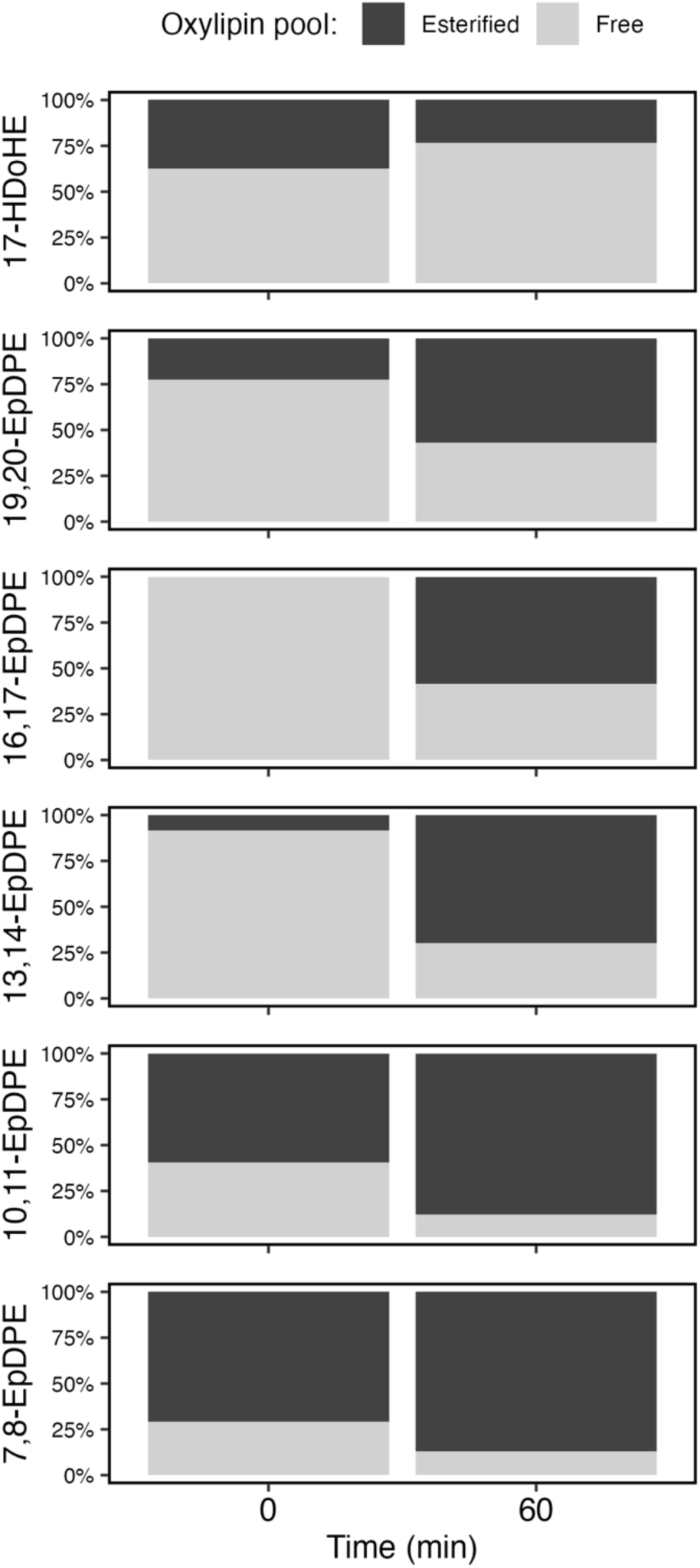
Percentage of free and calculated esterified (i.e., total - free) DHA-derived oxylipins from MFG-enriched fraction incubated at room temperature over 1 h in the presence of exogenous DHA (150 µM). Results are reported as the average of n=4. DHA: docosahexaenoic acid; HDoHE: hydroxydocosahexaenoic acid; EpDPE: epoxydocosahexaenoic acid.

### 3.3. MFG-enriched fraction becomes highly enriched in DHA-derived oxylipins upon incubation with DHA

To contextualize the contribution of DHA-derived oxylipins relative to those derived from other fatty acids, their cumulative percentages are shown in **Figure 3**. Upon incubation with DHA, the sum of all free oxylipins in the MFG-enriched fraction was 8.88 ± 1.34 pmol/mg of cream, of which DHA-derived species accounted for 7.27 ± 1.21 pmol/mg of cream—representing the majority (82.3 ± 9.6 %) of all free oxylipins in these samples. Conversely, the concentration of all free oxylipins in control samples (without exogenous DHA) was 0.49 ± 0.27 pmol/mg of cream, from which only 0.04 ± 0.05 pmol/mg cream (7.9 ± 8.9 %) derived from DHA. A similar trend was observed for total oxylipins. Their combined concentration in MFG-enriched fraction after incubation with DHA was 199.1 ± 36.4 pmol/mg cream, including 13.4 ± 5.3 pmol/mg of cream derived from DHA, representing 6.6 ± 1.9 %. In control samples (without exogenous DHA) the concentration of all total oxylipins was 125.7 ± 49.9 pmol/mg cream, but only 0.30 ± 0.07 pmol/mg cream (0.27 ± 0.08 %) from DHA. Overall, incubating the MFG-enriched fraction with DHA resulted in a 15 and 28-fold increment in the concentration of all free and total DHA-derived oxylipins, compared to the control samples without exogenous DHA.

**Figure 3.**
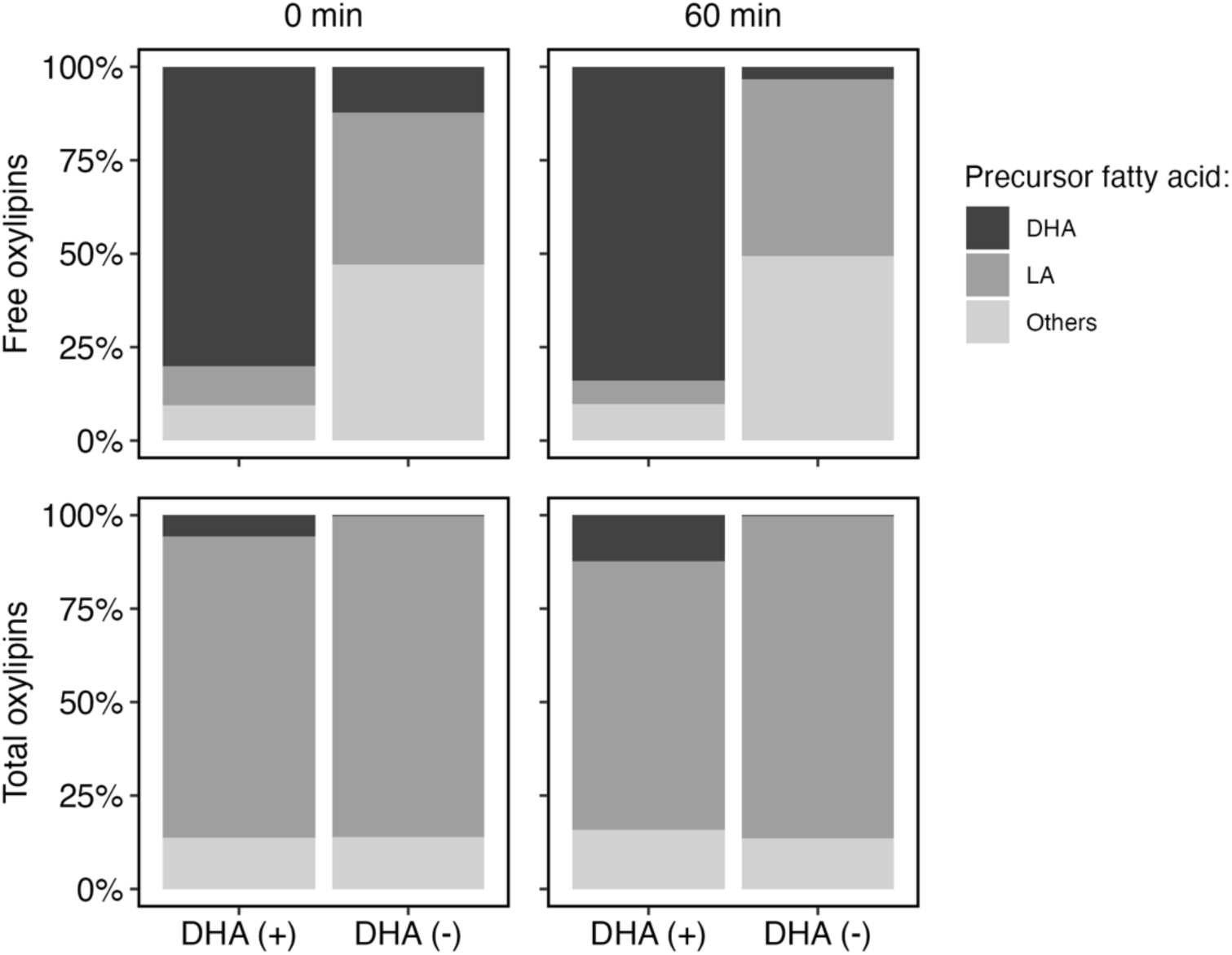
Oxylipin distribution in MFG-enriched fraction incubated over 1 h in the presence (+) or absence (-) of exogenous DHA (150 µM). Percentages are displayed based on precursor polyunsaturated fatty acids. Results are reported as the average of n=4. ‘Others’ refers to the cumulative of arachidonic acid, eicosapentaenoic acid, alfa linolenic acid, gamma-linolenic acid, and dihommo-gamma linolenic. DHA: docosahexaenoic acid; LA: linoleic acid.

### 3.4. Generation of oxylipins in MFG-enriched fraction upon incubation with exogenous DHA depends on MFG integrity

To assess the role of globule integrity in oxylipin generation in the MFG-enriched fraction following incubation with exogenous DHA, two approaches were employed. The first aimed to mechanically disrupt the MFGs by vortexing the samples during incubation; the second aimed to thermally treat the MFGs by preheating the MFG-enriched fraction at 90 °C for 10 minutes before the incubation with DHA. To visualize the impact of these treatments on MFG structure, fluorescent micrographs were acquired post-incubation, wherein the neutral lipids were stained with Nile Red (**Figure 4**).

**Figure 4.**
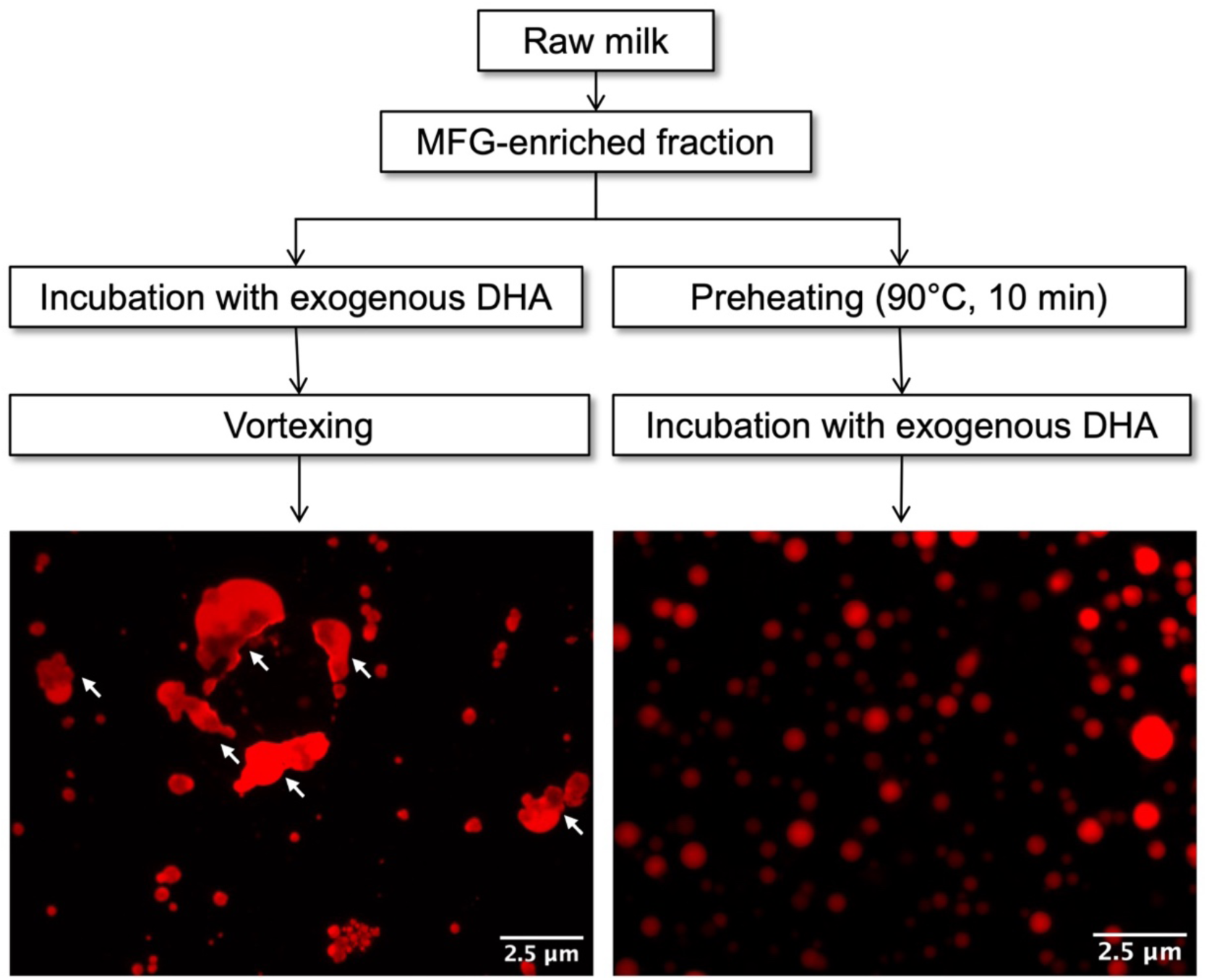
Effect of vortexing and preheating on the morphology of milk fat globules (MFG) from a MFG-enriched fraction. Neutral lipids in the MFG core were probed with Nile Red (red). Arrows highlight neutral lipids released from disrupted MFGs.

In the vortexed samples, neutral lipids were observed as both circular and non-circular, irregularly shaped structures, which indicates that while some MFGs were preserved, others were disrupted, leading to the release of neutral lipids from the core of the globules. In the preheated samples, only circular structures were observed, exhibiting preserved globular structures.

The concentrations of DHA-derived oxylipins in vortexed and preheated MFG-enriched fractions following incubation with or without exogenous DHA are shown in **Table 2** and **Table 3**, respectively. The concentrations of oxylipins derived from other PUFAs besides DHA in vortexed and preheated samples are reported in Supplementary Table S5 and Table S6.

**Table 2.** Effect of vortexing on the concentration of DHA-derived oxylipins (pmol/mg of cream) in a MFG-enriched fraction incubated at room temperature during 1h in the presence (+) or absence (-) of exogenous DHA (150 µM). Results are reported as the average ± standard deviation (n=3).

| Free Oxylipins |  | DHA (+) |  | DHA (-) |  |
| --- | --- | --- | --- | --- | --- |
|  |  | 0 min | 60 min | 0 min | 60 min |
| Epoxide | 19(20)-EpDPE | 0.7037 ± 0.1198 | 0.6628 ± 0.1424 | 0.1404 ± 0.2393 <sup>2</sup> | ND |
|  | 16(17)-EpDPE | 0.1613 ± 0.0226 | 0.1218 ± 0.0383 | 0.0316 ± 0.0511 <sup>2</sup> | 0.0638 ± 0.0117 |
|  | 13(14)-EpDPE | 0.1550 ± 0.0489 | 0.1313 ± 0.0162 | 0.0297 ± 0.0507 <sup>2</sup> | ND |
|  | 10(11)-EpDPE | 0.1863 ± 0.0720 | 0.1572 ± 0.0691 | 0.0338 ± 0.0562 <sup>2</sup> | ND |
|  | 7(8)-EpDPE | 0.1305 ± 0.0867 | 0.1101 ± 0.0354 | 0.0398 ± 0.0660 <sup>2</sup> | ND |
| Monohydroxy | 17-HDoHE | 1.5890 ± 0.5516 | 1.8286 ± 0.3502 | 0.3280 ± 0.5504 <sup>2</sup> | 0.0085 ± 0.0014 |

| Total oxylipins |  | DHA (+) |  | DHA (-) |  | DHA effect<br>p-value* |
| --- | --- | --- | --- | --- | --- | --- |
|  |  | 0 min | 60 min | 0 min | 60 min |  |
| Epoxide | 19(20)-EpDPE | 1.7771 ± 0.2134 | 3.5326 ± 0.9430 | 1.6200 ± 0.9999 | 1.1299 ± 0.2964 | 0.021 |
|  | 16(17)-EpDPE | 0.4978 ± 0.0296 | 0.2760 ± 0.0860 | 0.4214 ± 0.4484 | 0.0004 ± 0.0001 | 0.341 |
|  | 13(14)-EpDPE | 0.4170 ± 0.0346 | 0.3590 ± 0.0210 | 0.6200 ± 0.6936 | 0.2507 ± 0.0880 | 0.899 |
|  | 10(11)-EpDPE | 0.5751 ± 0.0230 | 0.3411 ± 0.0991 | 0.6362 ± 0.5489 | 0.2878 ± 0.1104 | 0.645 |
|  | 7(8)-EpDPE | 0.8622 ± 0.2268 | 0.6294 ± 0.1688 | 0.5319 ± 0.4441 | 0.4959 ± 0.1139 | 0.237 |
| Monohydroxy | 17-HDoHE | 2.4545 ± 0.9596 | 3.3734 ± 0.7952 | 0.9722 ± 0.5882 | 1.2191 ± 0.5166 | 0.001 |

\*From a two-way ANOVA with repeated measurements
Superscript number indicates number of imputed samples; imputation was performed as LOD/√2.
ND: non detected (compounds that were not found in 3 replicates per treatment).
MFG: milk fat globule; DHA: docosahexaenoic acid; EpDPE: epoxydocosapentaenoic acid; HDoHE: hydroxydocosahexaenoic acid.

**Table 3.** Effect of preheating on the concentration of free DHA-derived oxylipins (pmol/mg of cream) in a MFG-enriched fraction incubated at room temperature during 1h in the presence (+) or absence (-) of exogenous DHA (150 µM). Results are reported as the average ± standard deviation (n=4).

| Free Oxylipins |  | DHA (+) |  | DHA (-) |  |
| --- | --- | --- | --- | --- | --- |
|  |  | 0 min | 60 min | 0 min | 60 min |
| Epoxide | 19(20)-EpDPE | 2.7818 ± 1.5779 | 2.6142 ± 0.4943 | ND | ND |
| | 16(17)-EpDPE | 1.5335 ± 0.9409 | 1.3220 ± 0.3978 | ND | ND |
| | 13(14)-EpDPE | 1.3030 ± 0.7650 | 1.0664 ± 0.3842 | ND | 0.0008 ± 0.0007 <sup>2</sup> |
| | 10(11)-EpDPE | 0.7894 ± 0.2962 | 0.7236 ± 0.3606 | ND | ND |
| | 7(8)-EpDPE | 0.5406 ± 0.1427 | 0.6175 ± 0.2699 | 0.0011 ± 0.0007 <sup>2</sup> | 0.0030 ± 0.0025 <sup>2</sup> |
| Monohydroxy | 17-HDoHE | 0.7447 ± 0.3182 | 2.6095 ± 1.0968 | 0.0253 ± 0.0183 | 0.0241 ± 0.0232 <sup>2</sup> |

| Total oxylipins |  | DHA (+) |  | DHA (-) |  | DHA effect<br>p-value* |
| --- | --- | --- | --- | --- | --- | --- |
|  |  | 0 min | 60 min | 0 min | 60 min |  |
| Epoxide | 19(20)-EpDPE | 4.2197 ± 2.8069 | 2.9765 ± 0.9978 | 0.0856 ± 0.0437 | 0.0556 ± 0.0299 | 3.71 × 10<sup>-4</sup> |
| | 16(17)-EpDPE | 1.4063 ± 1.1122 | 0.8690 ± 0.2062 | 0.0153 ± 0.0165 <sup>1</sup> | 0.0183 ± 0.0178 <sup>1</sup> | 0.006 |
| | 13(14)-EpDPE | 1.6379 ± 1.3645 | 0.9651 ± 0.3363 | 0.0316 ± 0.0252 | 0.0155 ± 0.0046 | 0.003 |
| | 10(11)-EpDPE | 2.5153 ± 2.2802 | 1.2088 ± 0.6013 | 0.0180 ± 0.0125 | 0.0191 ± 0.0124 | 0.005 |
| | 7(8)-EpDPE | 2.3986 ± 2.0907 | 1.3196 ± 0.6270 | 0.0778 ± 0.0778 | 0.0265 ± 0.0122 | 0.005 |
| Monohydroxy | 17-HDoHE | 2.8732 ± 0.7795 | 2.9874 ± 0.5843 | 0.1679 ± 0.0708 | 0.1914 ± 0.0517 | 3.63 × 10<sup>-6</sup> |

\*From a two-way ANOVA with repeated measurements
Superscript number indicates number of imputed samples; imputation was performed as LOD/ $\sqrt{2}$ .
ND: non detected (compounds that were not found in 3-4 replicates per treatment).
MFG: milk fat globule; DHA: docosahexaenoic acid; EpDPE: epoxydocosapentaenoic acid; HDoHE: hydroxydocosahexaenoic acid.

In vortexed samples, when the MFG-enriched fraction was incubated with DHA, the concentrations of free 19(20)-EpDPE and 17-HDoHE (the most abundant DHA-oxylipins) were ∼0.7 and ∼1.7 pmol/mg cream, respectively. Meanwhile, in control samples without incubation with exogeneous DHA, the concentrations of free 19(20)-EpDPE and 17-HDoHE were lower, below 0.15 and 0.35 pmol/mg cream, respectively. ANOVA could not be performed for free DHA-oxylipins due to all the non-detected values in samples without addition of DHA.

In the same vortexed samples, two-way repeated measures ANOVA revealed that the addition of DHA significantly affected only the concentration of total 19-EpDPE and 17-HDoHE, wherein the increase in concentration following incubation with exogenous DHA was modest (less than 1 order of magnitude). Specifically, total 19(20)-EpDPE and 17-HDoHE levels ranged between 1.7 to 3.5 and 2.4 to 3.4 pmol/mg of cream, respectively, whereas in control samples, the concentration did not exceed 1.6 and 1.2 pmol/mg of cream. Although DHA-oxylipin concentrations were elevated in vortexed MFG-enriched samples upon incubation with DHA relative to control samples, this difference was substantially smaller than that observed in non-vortexed MFG-enriched samples, where oxylipin levels were 1–2 orders of magnitude higher than controls. These results suggest that vortexing, which causes significant structural disruption of MFGs, impairs the generation of both free and esterified DHA-derived oxylipin.

Incubation of the preheated MFG-enriched fraction with DHA resulted in significantly higher concentrations of DHA-oxylipins relative to control samples. In samples supplemented with DHA, concentration of free 19(20)-EpDPE and 17-HDoHE (here, too, the most abundant DHA-oxylipins) ranged from 2.6 to 2.8 and 0.7 to 2.6 pmol/mg cream, respectively. Meanwhile, in their control samples without exogenous DHA incubations these free oxylipins were undetected or below 0.02 pmol/mg cream. As with the other treatments, ANOVA could not be performed on free oxylipins because of the non-detected values in control samples.

Two-way repeated measures ANOVA confirmed that addition of DHA significantly affected the concentration of total DHA-oxylipins (*p*-value < 0.05) in the preheated samples. Total EpDPE and HDoHE levels in DHA-supplemented samples reached 0.8 to 4.2 and approximately 2.9 pmol/mg cream, respectively, representing 1–2 orders of magnitude higher levels than controls. These concentrations were comparable to those observed in non-preheated samples **Table 1**, indicating that preheating did not inhibit the generation of DHA-oxylipins. However, preheated samples showed no increase in total oxylipin concentration over time, suggesting that thermal treatment may have impaired the esterification of DHA-oxylipins

## 4. Discussion

The main finding of this study is that both free and esterified DHA-derived oxylipins were generated in a MFG-enriched fraction following incubation with exogenous DHA. These oxylipins included epoxide (19(20)-EpDPE, 16(17)-EpDPE, 13(14)-EpDPE, 10(11)-EpDPE, and 7(8)-EpDPE) and mono-hydroxy (17-HDoHE) derivatives of DHA.

In biological systems, oxylipins exist in the free and esterified forms (i.e., bound to neutral lipids and phospholipids). Free oxylipins are bioactive *in vivo* whereas the esterified pool is considered to play a major role in regulating free oxylipin availability and turnover.^18,47,48^ Particularly in milk, over 90% of oxylipins are esterified,^28^ and in both the free and esterified pools most oxylipins derive from linoleic acid, while DHA-oxylipins represent only 0.01-4.4% of the total ^27,28^, consistent with our control MFG-enriched samples. In contrast, incubation with exogenous DHA resulted in the MFG-enriched fraction becoming highly enriched in free and esterified DHA oxylipins—85 and 6%, respectively. In both pools, 17-HDoHE and 19(20)-EpDPE were the most abundant metabolites of DHA produced, reaching 2.1-3.3 and 2.2-5.1 pmol/mg, respectively. To put this concentration into context, ∼100 mg of MFG-enriched fraction after incubation with DHA displays an equivalent amount of these two oxylipins as 1 µL of fish oil, which contains ∼100-300 and ∼5-100 pmol/µL of 19(20)-EpDPE and 17-HDoHE, respectively ^59^. Thus, the MFG-enriched fraction incubated with DHA, as presented here, could represent a dietary approach to deliver bioactive lipids such as 19(20)-EpDPE and 17-HDoHE. The former exerts anti-hypertensive ^60^, anti-thrombotic ^61^, vasodilatory ^62^, and anti-angiogenic activities ^63^, while both are associated with antinociceptive properties ^64–66^. Future studies should evaluate whether the oxylipins produced in the MFG-enriched fraction are bound to neutral lipids or phospholipids, and overall, the bioavailability of these free and esterified oxylipins delivered via MFGs.

In mammals, epoxidation and hydroxylation of DHA, as well as esterification and hydrolysis of DHA oxylipins is a process catalyzed by membrane-bound enzymes, including CYP450, LOX, acyl-CoA synthetases and acyltransferases.^34,52,53^ Thus, we hypothesized that if the synthesis and esterification of oxylipins was enzymatic, disrupting the MFGs through mechanical or thermal means might reduce this activity. Indeed, vortexing led to significant damage to the MFGs and release of core neutral lipids, along with a marked reduction in the generation of both free and esterified DHA-oxylipins. This suggests that membrane integrity is essential for oxylipin synthesis in the MFG-enriched fraction, in agreement with the enzymatic hypothesis. Moreover, if the observed oxylipin production was non-enzymatic (i.e., autooxidation), vortexing would have enhanced oxylipin formation due to increased oxygen exposure, but this did not occur. Meanwhile, preheating of the MFG-enriched fraction prior to incubation with exogenous DHA did not impact generation levels of total DHA-oxylipins but appeared to affect their esterification. This result does not necessarily contradict an enzymatic pathway, as the putative responsible enzymes may have retained activity due to their thermal stability within the membrane under our experimental conditions. Notably, more than 90% of xanthine oxidoreductase activity (a known MFGM-associated enzyme) can be retained following batch pasteurization or high-temperature short-time milk processing ^22^. Furthermore, in our experiments the MFG-enriched fraction (∼40% w/v fat ^30^, similar to milk cream) was diluted to 20% w/v in PBS before heating. Higher fat concentrations have been associated with reduced release of xanthine oxidoreductase from the MFGM ^54^, suggesting greater membrane stability under these conditions.

Although oxylipins can indeed be formed through the autooxidation of DHA, we consider this mechanism unlikely to account for our findings, for two reasons. First, no increase in DHA-derived oxylipins was observed in DHA stock incubated without the MFG-enriched fraction under identical conditions (Supplementary Table S4). Second, the non-enzymatic generation of these compounds typically occurs over several days,^55–58^ whereas the changes observed in this study occurred within minutes. Furthermore, technical artifacts during solid phase extraction of oxylipins that can lead to autooxidation were not expected here, given the use of Oasis-HLB columns, which have been shown to minimize autooxidation.^59^

We thus propose that the data presented here, demonstrating that free and esterified EpDPE and HDoHE were generated in an MFG-enriched fraction upon incubation with DHA, may be explained by the endogenous presence of CYP450, LOX, acyl-CoA synthetase, acyltransferases and lipases within the MFGs. Given that the inner membrane of the MFGM originates from the endoplasmic reticulum of mammary epithelial cells, it is plausible that trace amounts of these ER membrane-bound enzymes are incorporated into the MFGM during secretion, as it occurs with other known MFGM-bound enzymes such as xanthine oxidoreductase (reviewed by ^20,23,24^).

Several studies suggest the presence of CYP450, LOX, acyl-CoA synthetase, acyltransferases and lipases within the MFGM or mammary epithelial tissue (**Table *4***). For instance, in the 1970’s, researchers reported the presence of the inactive form of CYP450 and a b-type cytochrome (an electron donor for CYP450 activity ^33^) in MFGM ^34,35^ isolated from bovine milk. Based on the detection limits of the analytical methods used, authors estimated that if active CYP450 was present, its levels must have been <10 pmol/mg protein (less than 5% of what typically found in liver microsomes) ^34^. More recently, proteomic studies have identified CYP450, cytochrome b5, and arachidonate 15 and 12-LOX in the

**Table 4.**
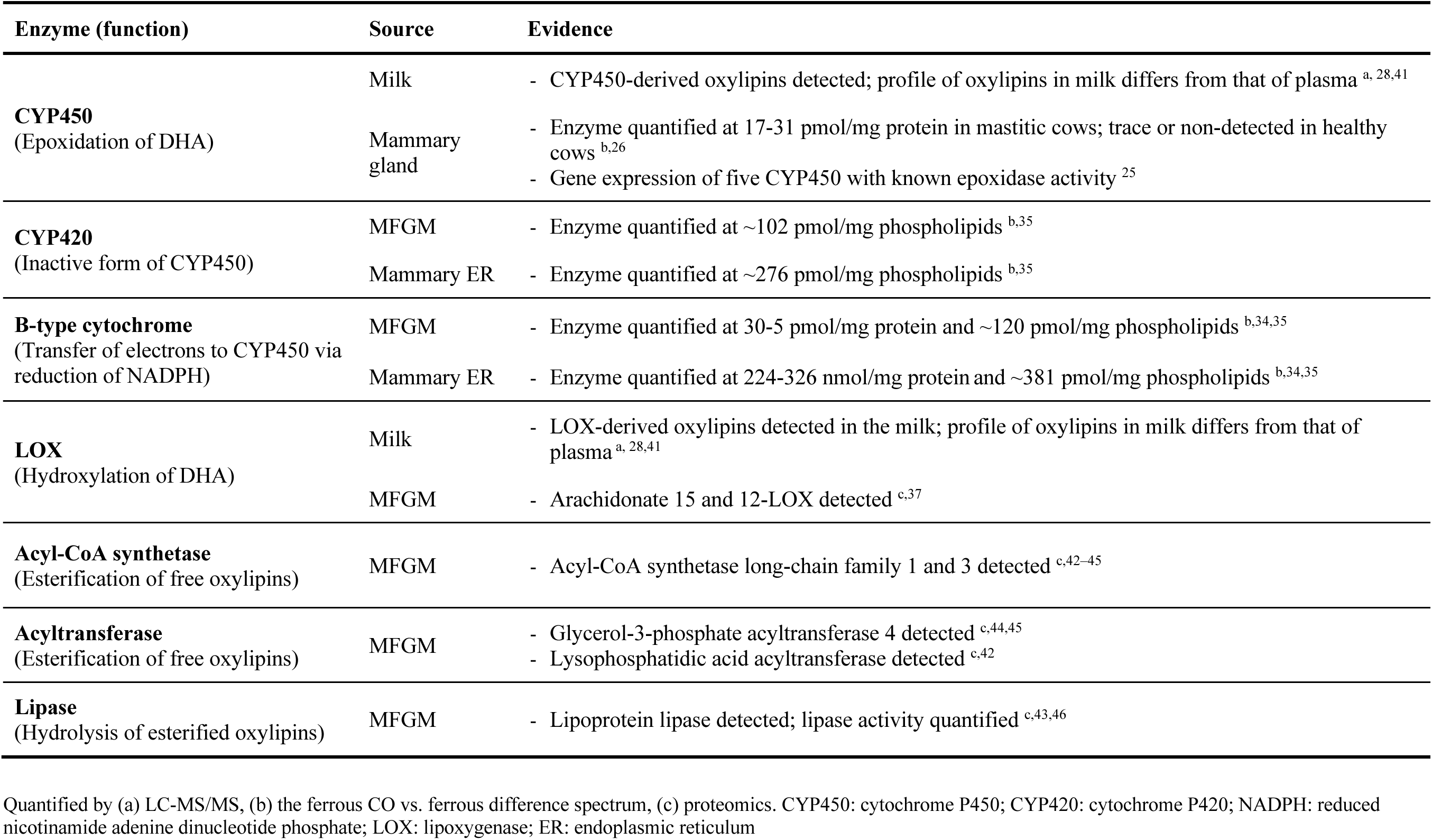
Evidence of the presence of enzymes involved in the transformation of docosahexaenoic acid (DHA) to epoxydocosapentaenoic acid (EpDPE) and hydroxydocosahexaenoic acid (HDoHE) in bovine milk, milk fat globule membrane (MFGM) or mammary gland tissue.

MFGM of human, bovine, goat, and yak milk ^36–39^. Meanwhile, in bovine mammary epithelial tissue (from mastitic cows), CYP450 concentrations of 17–31 pmol/mg protein were quantified ^26^. The expression of five CYP450 isoforms with known epoxygenase activity were detected in the mammary tissue of healthy lactating cows: CYP2C19, CYP1A1, CYP2E1, CYP3A4, and CYP2J2 ^25^. These isoforms exhibit DHA epoxidation rates of 1.6, 0.5, 0.8, 0.7, and 0.2 pmol/min/pmol CYP450, respectively ^40^. Except for CYP2C19, these enzymes preferentially epoxidize the last double bond of DHA, predominantly forming 19(20)-EpDPE ^40^— the most abundant DHA epoxide observed in this study. Acyl-CoA synthetase, acyltransferases and lipases have all been reported in the MFGM ^42–46^, and the lipase activity of this fraction has been shown before ^43,46^ (**Table *4***).

In summary, the detection of free and esterified EpDPEs and HDoHE in the MFG-enriched fraction following DHA incubation strongly supports the presence of multiple active enzymes. We thus propose a mechanism that would explain these observations (Figure 5). First, exogenous DHA in free form partitions into the MFG where it is enzymatically oxidized to epoxides and mono-hydroxy oxylipins by the action of endogenous CYP450 and LOX enzymes, respectively. Second, free oxylipins could then be esterified to neutral lipids or phospholipids by acyl-CoA synthetase and acyltransferases, while esterified oxylipins could be released from their bound state by lipases. We also propose that the rate of esterification outpaced both oxylipin synthesis from DHA and/or hydrolysis of esterified oxylipins, as we observed that during the 60-minute incubation period the esterified/free oxylipin ratio increased despite stable concentrations of free oxylipins. A key limitation of this study is that the presence of these enzymes was not directly confirmed, because the specific isoforms involved are yet to be characterized in bovine milk. Future studies should address this gap by employing specific enzymatic inhibitors, immunodetection techniques, or enzyme quantification methods to identify and quantify these putative enzymes. The latter would allow for more precise estimation of DHA epoxidation and hydroxylation rates normalized to enzyme concentration.

**Figure 5.**
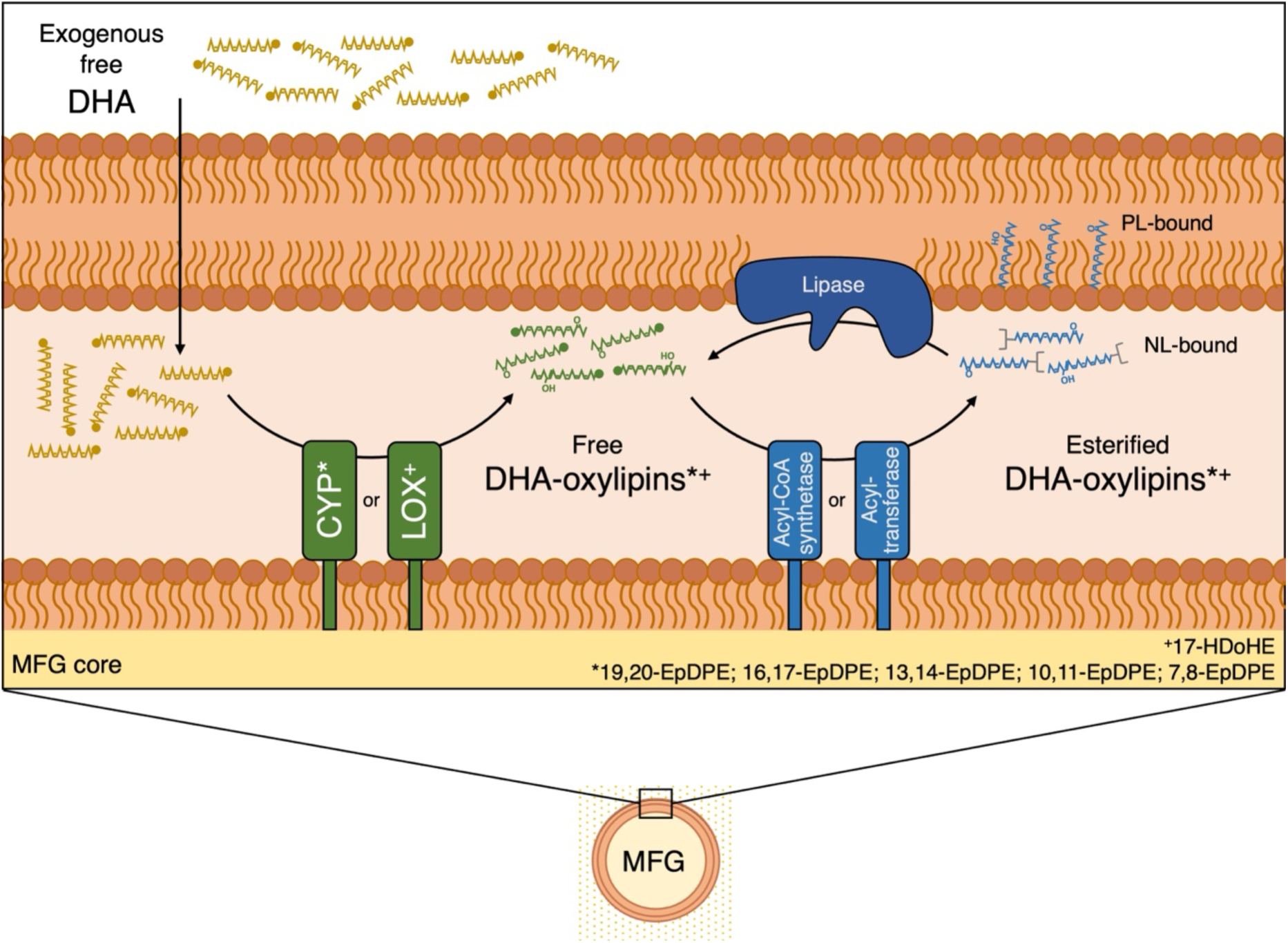
Proposed mechanism explaining the generation of DHA-oxylipins in a milk fat globule (MFG)-enriched fraction following incubation with exogenous docosahexaenoic acid (DHA). DHA partitions into the MFG where it is enzymatically oxidized to free oxylipins by the action of cytochrome p450 (CYP) and lipoxygenases (LOX). Free oxylipins are then esterified to phospholipids (PL) or neutral lipids (NL) by acyl-CoA synthetases or acyltransferases or hydrolyzed from them by lipases. The location of the enzymes is speculative. EpDPE: epoxydocosapentaenoic acid; HDoHE: hydroxydocosahexaenoic acid.

Other food ingredients like soybean and oat flour have been shown to retain their LOX and peroxygenase activity, and to have the capacity to catalyze the conversion of EPA and DHA to their epoxide, hydroxy and di-hydroxy oxylipins ^67,68^. While the MFG-enriched fraction might yield lower rates of PUFA to oxylipin conversion than what was observed using these enzyme-rich flours or microbial enzymes ^16,69,70^, the MFG-enriched fraction might be more suitable for certain applications such as infant nutrition. For instance, oxylipins in milk are thought to regulate neonatal development and immunity.^71–73^ Moreover, while the natural presence of oxylipins in milk is attributed to their production by mammary epithelial cells ^25,41,74^, we propose that this biocatalytic potential is inherited by the MFG, which raises the question about the possible biological significance of carrying these enzymes within the MFGM.

## 5. Conclusion

This study presents the first evidence that a MFG-enriched fraction (isolated from raw and unhomogenized bovine milk) display the biocatalytic potential to transform exogenous DHA into its oxylipins. This biocatalytic activity may extend to the production of other clinically and nutritionally relevant compounds, including oxylipins derived from other PUFAs. The consumption of the MFG-enriched fraction following incubation with DHA, as presented here, could represent an approach to deliver bioactive lipids, provided that the bioavailability of dietary DHA-derived oxylipins is shown in future studies. Overall, our findings support the dual use of MFGs as food-grade biocatalysts and encapsulation matrices for the synthesis and delivery of bioactive lipids and suggests that MFGs contain various metabolically active enzymes that can work cooperatively, such as CYP450, LOX, and acyltransferases/acyl-CoA synthetases.

## 6. Declarations

### 6.2 Acknowledgments

The authors acknowledge Mayuko Itaya, from Tohoku University, for her initial contributions to this project.

### 6.2. Contributions

T.H.B. designed and performed all experiments, analyzed the data, and wrote the manuscript. N.N. and A.Y.T. reviewed and edited the manuscript, supervised the work and secured the funding.

## 7. Supplementary material

Table S1 – Analytical parameters for the 76 oxylipins measured by UPLC-MS/MS

Table S2 – *p*-values of statistical analyses performed.

Table S3 – Concentration of oxylipins from a MFG-enriched fraction incubated with or without exogenous DHA.

Table S4 – Concentration of DHA-oxylipins in a DHA stock.

Table S5 – Concentration of oxylipins from a preheated MFG-enriched fraction incubated with or without exogenous DHA.

Table S6 – Concentration of DHA-oxylipins from a vortexed MFG-enriched fraction incubated with or without exogenous DHA.

## Bibliography

1. Dyall, S. C. et al. Polyunsaturated fatty acids and fatty acid-derived lipid mediators: Recent advances in the understanding of their biosynthesis, structures, and functions. Prog. Lipid Res. 86, 101165 (2022).

2. Borsini, A. et al. Omega-3 polyunsaturated fatty acids protect against inflammation through production of LOX and CYP450 lipid mediators: relevance for major depression and for human hippocampal neurogenesis. Mol. Psychiatry 26, 6773–6788 (2021).

3. Yang, Y. et al. Differential Effects of 17,18-EEQ and 19,20-EDP Combined with Soluble Epoxide Hydrolase Inhibitor t-TUCB on Diet-Induced Obesity in Mice. Int. J. Mol. Sci. 22, (2021).

4. Ontko, C. D., Capozzi, M. E., Kim, M. J., McCollum, G. W. & Penn, J. S. Cytochrome P450-epoxygenated fatty acids inhibit Müller glial inflammation. Sci. Rep. 11, 9677 (2021).

5. Xiao, Y. et al. Bioactive oxylipins in type 2 diabetes mellitus patients with and without hypertriglyceridemia. Front. Endocrinol. (Lausanne) 14, 1195247 (2023).

6. Hateley, C. et al. Multi-tissue profiling of oxylipins reveal a conserved up-regulation of epoxide:diol ratio that associates with white adipose tissue inflammation and liver steatosis in obesity. EBioMedicine 103, 105127 (2024).

7. Chiang, K.-M. et al. Identification of Serum Oxylipins Associated with the Development of Coronary Artery Disease: A Nested Case-Control Study. Metabolites 12, (2022).

8. Ostermann, A. I. et al. Plasma oxylipins respond in a linear dose-response manner with increased intake of EPA and DHA: results from a randomized controlled trial in healthy humans. Am. J. Clin. Nutr. 109, 1251–1263 (2019).

9. Gabbs, M., Zahradka, P., Taylor, C. G. & Aukema, H. M. Time Course and Sex Effects of α-Linolenic Acid-Rich and DHA-Rich Supplements on Human Plasma Oxylipins: A Randomized Double-Blind Crossover Trial. J. Nutr. 151, 513–522 (2021).

10. Downie, C. G. et al. Genome-wide association study reveals shared and distinct genetic architecture of fatty acids and oxylipins in the Hispanic Community Health Study/Study of Latinos. HGG Adv. 6, 100390 (2025).

11. Saleh, R. N. M. et al. APOE Genotype Modifies the Plasma Oxylipin Response to Omega-3 Polyunsaturated Fatty Acid Supplementation in Healthy Individuals. Front. Nutr. 8, 723813 (2021).

12. Hwang, S. H. et al. Chemical synthesis and biological evaluation of ω-hydroxy polyunsaturated fatty acids. Bioorg. Med. Chem. Lett. 27, 620–625 (2017).

13. Cinelli, M. A. & Lee, K. S. S. Asymmetric Total Synthesis of 19,20-Epoxydocosapentaenoic Acid, a Bioactive Metabolite of Docosahexaenoic Acid. J. Org. Chem. 84, 15362–15372 (2019).

14. Nanba, Y., Shinohara, R., Morita, M. & Kobayashi, Y. Stereoselective synthesis of 17,18-epoxy derivative of EPA and stereoisomers of isoleukotoxin diol by ring opening of TMS-substituted epoxide with dimsyl sodium. Org. Biomol. Chem. 15, 8614–8626 (2017).

15. Rorrer, G. L. et al. Bioreactor seaweed cell culture for production of bioactive oxylipins. J Appl Phycol 7, 187–198 (1995).

16. Woodman, J. W., Cinelli, M. A., Scharmen-Burgdolf, A. & Lee, K. S. S. Enzymatic synthesis of epoxidized metabolites of docosahexaenoic, eicosapentaenoic, and arachidonic acids. J. Vis. Exp. (2019). doi:10.3791/59770

17. González-Benjumea, A. et al. Regioselective and Stereoselective Epoxidation of n-3 and n-6 Fatty Acids by Fungal Peroxygenases. Antioxidants (Basel) 10, (2021).

18. Zhang, Z. et al. Linoleic acid-derived 13-hydroxyoctadecadienoic acid is absorbed and incorporated into rat tissues. Biochim. Biophys. Acta Mol. Cell Biol. Lipids 1866, 158870 (2021).

19. Moser, B. R., Cermak, S. C., Doll, K. M., Kenar, J. A. & Sharma, B. K. A review of fatty epoxide ring opening reactions: Chemistry, recent advances, and applications. J Am Oil Chem Soc 99, 801–842 (2022).

20. Keenan, T. W. & Mather, I. H. in Advanced dairy chemistry volume 2 lipids (eds. Fox, P. F. & McSweeney, P. L. H.) 137–171 (Springer US, 2006). doi:10.1007/0-387-28813-9_4

21. Ozturk, G., Shah, I. M., Mills, D. A., German, J. B. & de Moura Bell, J. M. L. N. The antimicrobial activity of bovine milk xanthine oxidase. Int. Dairy J. 102, (2020).

22. Ozturk, G., German, J. B. & de Moura Bell, J. M. L. N. Effects of industrial heat treatments on the kinetics of inactivation of antimicrobial bovine milk xanthine oxidase. *npj Sci*. Food 3, 13 (2019).

23. Patton, S. Origin of the milk fat globule. J Am Oil Chem Soc 50, 178–185 (1973).

24. Wooding, F. B. P. & Kinoshita, M. Milk fat globule membrane: formation and transformation. J. Dairy Res. 90, 367–375 (2023).

25. Kuhn, M. J., Putman, A. K. & Sordillo, L. M. Widespread basal cytochrome P450 expression in extrahepatic bovine tissues and isolated cells. J. Dairy Sci. 103, 625–637 (2020).

26. Atroshi, F., Kaipainen, P. & Parantainen, J. Evidence for the presence of cytochrome P-450 in mastitic bovine mammary gland. Pharmacol. Res. Commun. 19, 673–678 (1987).

27. Gan, J., Zhang, Z., Kurudimov, K., German, J. B. & Taha, A. Y. Distribution of free and esterified oxylipins in cream, cell, and skim fractions of human milk. Lipids 55, 661–670 (2020).

28. Teixeira, B. F., Dias, F. F. G., Vieira, T. M. F. de S., Leite Nobrega de Moura Bell, J. M. & Taha, A. Y. Method optimization of oxylipin hydrolysis in nonprocessed bovine milk indicates that the majority of oxylipins are esterified. J. Food Sci. 86, 1791–1801 (2021).

29. Alshehab, M., Hernández-Barrueta, T., Cui, H. & Nitin, N. Milk fat globules—A natural carrier for exogenous lipophilic bioactives. Food Res. Int 116891 (2025). doi:10.1016/j.foodres.2025.116891

30. Alshehab, M., Reis, M. G., Day, L. & Nitin, N. Milk fat globules, a novel carrier for delivery of exogenous cholecalciferol. Food Res. Int. 126, 108579 (2019).

31. Gallier, S., Gragson, D., Jiménez-Flores, R. & Everett, D. Using confocal laser scanning microscopy to probe the milk fat globule membrane and associated proteins. J. Agric. Food Chem. 58, 4250–4257 (2010).

32. Greenspan, P., Mayer, E. P. & Fowler, S. D. Nile red: a selective fluorescent stain for intracellular lipid droplets. J. Cell Biol. 100, 965–973 (1985).

33. Henderson, C. J., McLaughlin, L. A. & Wolf, C. R. Evidence that cytochrome b5 and cytochrome b5 reductase can act as sole electron donors to the hepatic cytochrome P450 system. Mol. Pharmacol. 83, 1209–1217 (2013).

34. Jarasch, E. D., Bruder, G., Keenan, T. W. & Franke, W. W. Redox constituents in milk fat globule membranes and rough endoplasmic reticulum from lactating mammary gland. J. Cell Biol. 73, 223–241 (1977).

35. Bruder, G., Fink, A. & Jarasch, E. D. The B-type cytochrome in endoplasmic reticulum of mammary gland epithelium and milk fat globule membranes consists of two components cytochrome b5 and cytochrome P-420. Exp. Cell Res. 117, 207–217 (1978).

36. He, B. et al. Proteomics profiles of bovine and goat milk fat globule membrane revealed differences on the components and potential functions. Int. J. Dairy Technol. 78, (2025).

37. Ji, Z. et al. Comprehensive analysis of species-specific differences in fatty acid composition and proteome of milk fat globules in human and animals. Food Chemistry: X 27, 102431 (2025).

38. Wang, Y. et al. Astral-data-independent acquisition depicts the dynamic landscape of milk fat globule membrane proteins in yak colostrum, transitional milk, and mature milk. J. Dairy Sci. (2025). doi:10.3168/jds.2025-26661

39. Tang, J. et al. Effects of maternal and perinatal factors on human milk fat globule membrane proteome: A data independent acquisition approach. Food Biosci. 58, 103791 (2024).

40. Fer, M. et al. Metabolism of eicosapentaenoic and docosahexaenoic acids by recombinant human cytochromes P450. Arch. Biochem. Biophys. 471, 116–125 (2008).

41. Kuhn, M. J. et al. Differences in the oxylipid profiles of bovine milk and plasma at different stages of lactation. J. Agric. Food Chem. 65, 4980–4988 (2017).

42. Reinhardt, T. A. & Lippolis, J. D. Developmental changes in the milk fat globule membrane proteome during the transition from colostrum to milk. J. Dairy Sci. 91, 2307–2318 (2008).

43. Zhao, L., Du, M., Gao, J., Zhan, B. & Mao, X. Label-free quantitative proteomic analysis of milk fat globule membrane proteins of yak and cow and identification of proteins associated with glucose and lipid metabolism. Food Chem. 275, 59–68 (2019).

44. Lu, J., van Hooijdonk, T., Boeren, S., Vervoort, J. & Hettinga, K. Identification of lipid synthesis and secretion proteins in bovine milk. J. Dairy Res. 81, 65–72 (2014).

45. Lu, J. et al. The protein and lipid composition of the membrane of milk fat globules depends on their size. J. Dairy Sci. 99, 4726–4738 (2016).

46. Lu, J. et al. Comparative proteomics of milk fat globule membrane in different species reveals variations in lactation and nutrition. Food Chem. 196, 665–672 (2016).

47. Otoki, Y. et al. Acute hypercapnia/ischemia alters the esterification of arachidonic acid and docosahexaenoic acid epoxide metabolites in rat brain neutral lipids. Lipids 55, 7–22 (2020).

48. Watanabe, S. et al. Intraperitoneally injected d11-11(12)-epoxyeicosatrienoic acid is rapidly incorporated and esterified within rat plasma and peripheral tissues but not the brain. Prostaglandins Leukot Essent Fatty Acids 202, 102622 (2024).

49. Liu, G.-Y. et al. Synthesis of oxidized phospholipids by sn-1 acyltransferase using 2-15-HETE lysophospholipids. J. Biol. Chem. 294, 10146–10159 (2019).

50. Klett, E. L., Chen, S., Yechoor, A., Lih, F. B. & Coleman, R. A. Long-chain acyl-CoA synthetase isoforms differ in preferences for eicosanoid species and long-chain fatty acids. J. Lipid Res. 58, 884–894 (2017).

51. Shearer, G. C. & Newman, J. W. Lipoprotein lipase releases esterified oxylipins from very low-density lipoproteins. Prostaglandins Leukot Essent Fatty Acids 79, 215–222 (2008).

52. Lewin, T. M., Kim, J. H., Granger, D. A., Vance, J. E. & Coleman, R. A. Acyl-CoA synthetase isoforms 1, 4, and 5 are present in different subcellular membranes in rat liver and can be inhibited independently. J. Biol. Chem. 276, 24674–24679 (2001).

53. Shornick, L. P. & Holtzman, M. J. A cryptic, microsomal-type arachidonate 12-lipoxygenase is tonically inactivated by oxidation-reduction conditions in cultured epithelial cells. J. Biol. Chem. 268, 371–376 (1993).

54. Holzmüller, W., Müller, M., Himbert, D. & Kulozik, U. Impact of cream washing on fat globules and milk fat globule membrane proteins. International Dairy Journal 59, 52–61 (2016).

55. Reynaud, D., Thickitt, C. P. & Pace-Asciak, C. R. Facile preparation and structural determination of monohydroxy derivatives of docosahexaenoic acid (HDoHE) by alpha-tocopherol-directed autoxidation. Anal. Biochem. 214, 165–170 (1993).

56. Ferraz Teixeira, B., Furlan Gonçalves Dias, F., Ferreira de Souza Vieira, T. M., Y Taha, A. & Leite Nobrega de Moura Bell, J. M. Early detection of lipid oxidation in infant milk formula by measuring free oxylipins-Comparison with hydroperoxide value and thiobarbituric acid reactive substance methods. J. Food Sci. 87, 5252–5262 (2022).

57. Shen, Q. et al. Triacylglycerols are preferentially oxidized over free fatty acids in heated soybean oil. *npj Sci*. Food 5, 7 (2021).

58. Furlan Gonçalves Dias, F., Teixeira, B. F., Vieira, T. M. F. de S., de Moura Bell, J. M. L. N. & Taha, A. Y. Storage duration and added docosahexaenoic acid modify the rates of esterified and free oxylipin formation in infant milk formula. Processes 11, 3045 (2023).

59. Emami, S., Zhang, Z. & Taha, A. Y. Quantitation of Oxylipins in Fish and Algae Oil Supplements Using Optimized Hydrolysis Procedures and Ultra-High Performance Liquid Chromatography Coupled to Tandem Mass-Spectrometry. J. Agric. Food Chem. 68, 9329–9344 (2020).

60. Ulu, A. et al. Anti-inflammatory effects of ω-3 polyunsaturated fatty acids and soluble epoxide hydrolase inhibitors in angiotensin-II-dependent hypertension. J. Cardiovasc. Pharmacol. 62, 285–297 (2013).

61. Jung, F. et al. Effect of cytochrome P450-dependent epoxyeicosanoids on Ristocetin-induced thrombocyte aggregation. Clin Hemorheol Microcirc 52, 403–416 (2012).

62. Ye, D. et al. Cytochrome p-450 epoxygenase metabolites of docosahexaenoate potently dilate coronary arterioles by activating large-conductance calcium-activated potassium channels. J. Pharmacol. Exp. Ther. 303, 768–776 (2002).

63. Zhang, G. et al. Epoxy metabolites of docosahexaenoic acid (DHA) inhibit angiogenesis, tumor growth, and metastasis. Proc. Natl. Acad. Sci. USA 110, 6530–6535 (2013).

64. Valdes, A. M. et al. Association of the resolvin precursor 17-HDHA, but not D- or E- series resolvins, with heat pain sensitivity and osteoarthritis pain in humans. Sci. Rep. 7, 10748 (2017).

65. Ramsden, C. E. et al. Targeted alteration of dietary n-3 and n-6 fatty acids for the treatment of chronic headaches: a randomized trial. Pain 154, 2441–2451 (2013).

66. Morisseau, C. et al. Naturally occurring monoepoxides of eicosapentaenoic acid and docosahexaenoic acid are bioactive antihyperalgesic lipids. J. Lipid Res. 51, 3481–3490 (2010).

67. Sanfilippo, C., Paterna, A., Biondi, D. M. & Patti, A. Lyophilized extracts from vegetable flours as valuable alternatives to purified oxygenases for the synthesis of oxylipins. Bioorg Chem 93, 103325 (2019).

68. Tu, H.-A. T., Dobson, E. P., Henderson, L. C., Barrow, C. J. & Adcock, J. L. Soy flour as an alternative to purified lipoxygenase for the enzymatic synthesis of resolvin analogues. N. Biotechnol. 41, 25–33 (2018).

69. Speckmann, B. et al. Synbiotic Compositions of Bacillus megaterium and Polyunsaturated Fatty Acid Salt Enable Self-Sufficient Production of Specialized Pro-Resolving Mediators. Nutrients 14, (2022).

70. Shin, K.-C. et al. Enzymatic Formation of Protectin Dx and Its Production by Whole-Cell Reaction Using Recombinant Lipoxygenases. Catalysts 12, 1145 (2022).

71. Weiss, G. A. et al. High levels of anti-inflammatory and pro-resolving lipid mediators lipoxins and resolvins and declining docosahexaenoic acid levels in human milk during the first month of lactation. Lipids Health Dis. 12, 89 (2013).

72. Arnardottir, H., Orr, S. K., Dalli, J. & Serhan, C. N. Human milk proresolving mediators stimulate resolution of acute inflammation. Mucosal Immunol. 9, 757–766 (2016).

73. Alexandre-Gouabau, M.-C. et al. Breast Milk Lipidome Is Associated with Early Growth Trajectory in Preterm Infants. Nutrients 10, (2018).

74. King, K. et al. Dynamics of lipid droplets in the endometrium and fatty acids and oxylipins in the uterine lumen, blood, and milk of lactating cows during diestrus. J. Dairy Sci. 104, 3676–3692 (2021).

